# *shani* mutation in mouse affects splicing of Spata22 and leads to impaired meiotic recombination

**DOI:** 10.1101/2020.02.04.934042

**Authors:** Cynthia Petrillo, Vilma Barroca, Jonathan Ribeiro, Nathalie Lailler, Gabriel Livera, Scott Keeney, Emmanuelle Martini, Devanshi Jain

## Abstract

Recombination is crucial for chromosome pairing and segregation during meiosis. SPATA22, along with its direct binding partner and functional collaborator, MEIOB, is essential for the proper repair of double-strand breaks (DSBs) during meiotic recombination. Here we describe a novel point-mutated allele (*shani*) of mouse *Spata22* that we isolated in a forward genetic screen. *shani* mutant mice phenocopy *Spata22*-null and *Meiob*-null mice: mutant cells appear to form DSBs and initiate meiotic recombination, but are unable to complete DSB repair, leading to meiotic prophase arrest, apoptosis and sterility. *shani* mutants show precocious loss of DMC1 foci and improper accumulation of BLM-positive recombination foci, reinforcing the requirement of SPATA22-MEIOB for the proper progression of meiotic recombination events. The *shani* mutation lies within a *Spata22* coding exon and molecular characterization shows that it leads to incorrect splicing of the *Spata22* mRNA, ultimately resulting in no detectable SPATA22 protein. We propose that the *shani* mutation alters an exonic splicing enhancer element (ESE) within the *Spata22* transcript. The affected DNA nucleotide is conserved in most tetrapods examined, suggesting that the splicing regulation we describe here may be a conserved feature of *Spata22* regulation.

## Introduction

Gamete formation during sexual reproduction occurs by a specialized cell division program called meiosis, where two rounds of chromosome segregation follow one round of DNA replication; homologous chromosomes segregate in the first round and sister chromatids separate in the second. The meiotic cell division program contains an extended prophase, during which replicated homologous chromosomes pair and recombine. This creates temporary connections called crossovers that are essential to stabilize homologs on the metaphase spindle during segregation (Page and Hawley 2003; Hunter 2015). Errors in these events can cause gametogenic failure and infertility, or can produce aneuploid gametes that in turn result in miscarriage and developmental defects (Hassold and Hunt 2001; Sasaki et al. 2010).

Meiotic recombination initiates with the formation of programmed DSBs that are resected to yield single-strand DNA (ssDNA) overhangs. Then, DMC1, the meiosis-specific strand-exchange protein, along with its ubiquitously expressed paralog, RAD51, assembles onto ssDNA to form nucleoprotein filaments that conduct homology search and strand invasion. The resulting recombination intermediates are repaired by multiple pathways to ultimately give either non-crossover products or crossovers (Baudat et al. 2013; Hunter 2015). Recombination intermediates and their repair pathways are closely regulated to guarantee the formation of at least one crossover per homolog pair (Hunter 2015; Zickler and Kleckner 2015). While we have a broad-strokes picture of the factors involved and their contributions to meiotic recombination, mechanistic detail of their individual molecular functions and their cross-talk is lacking. We also lack clear understanding of how meiosis-specific proteins collaborate with ubiquitous repair proteins to specialize homologous recombination during meiosis. Moreover, it is unclear whether the catalog of relevant vertebrate proteins important for meiotic recombination is complete.

To explore the mechanisms underlying the regulation of meiotic recombination and to identify novel mouse meiotic mutants, we performed a phenotype-based, random chemical mutagenesis screen. Two hits we previously described are *rahu*, that was defective for a rodent-specific DNA methyltransferase paralog DNMT3C (Jain et al. 2017), and *ketu*, which was defective for the conserved RNA helicase YTHDC2 (Jain et al. 2018). Here, we describe a new mutant that we named *shani*, for ‘***<u>S</u>****pata22*-affected with **<u>h</u>**ypogonadic testes **<u>an</u>**d **i**nfertility’. Shani, Ketu and Rahu are harbingers of misfortune in Vedic mythology.

*shani* is a missense mutation in *Spata22*, which encodes a protein that is essential for meiotic recombination (La Salle et al. 2012; Ishishita et al. 2013). The *shani* mutation is expected to alter an amino acid, but surprisingly it causes exon skipping, eventually leading to lack of detectable SPATA22 protein. SPATA22 and its functional collaborator, MEIOB, form a meiosis-specific complex and they resemble subunits of RPA, the ubiquitous ssDNA-binding complex that regulates ssDNA during replication, repair and recombination (Wold 1997; Luo et al. 2013; Souquet et al. 2013; Ishishita et al. 2014; Ribeiro et al. 2016; Xu et al. 2017; Ribeiro et al. 2018). SPATA22-MEIOB colocalize on meiotic chromosomes with RPA subunits and their null mutants accumulate unrepaired DSBs, suggesting that they perform a critical function(s) during meiotic recombination (La Salle et al. 2012; Luo et al. 2013; Souquet et al. 2013; Ishishita et al. 2014; Hays et al. 2017). Also, mutations in this complex have been linked to human infertility (Gershoni et al. 2017; Caburet et al. 2019; Gershoni et al. 2019).

*Spata22* ^*shani/shani*^ homozygote mice are both female- and male-sterile. We show that *Spata22*^*shani/shani*^ mutant spermatocytes have defective chromosome synapsis and fail to complete DSB repair during meiotic prophase, like *Spata22*-null mice (Hays et al. 2017). Our results validate previous findings as well as further our understanding of the meiotic role of SPATA22 in mice. Importantly, we have isolated a novel allele of *Spata22* that reveals unexpected post-transcriptional regulation and may provide insight into how gene expression is regulated more generally during meiosis.

## Results

### Isolation of the novel meiotic mutant shani from a forward genetic screen

To isolate mutants with defects in meiosis, we carried out a phenotype based, random mutagenesis screen in mice (Jain et al. 2017; Jain et al. 2018). We mutagenized male mice with the alkylating agent *N*-ethyl-*N*-nitrosourea (ENU) that predominantly creates single base substitutions. Then we followed a three-generation breeding scheme to recover recessive mutations causing meiotic defects (Caspary 2010; Probst and Justice 2010; Jain et al. 2017; Jain et al. 2018)(**Fig. 1a**).

**Fig. 1.**
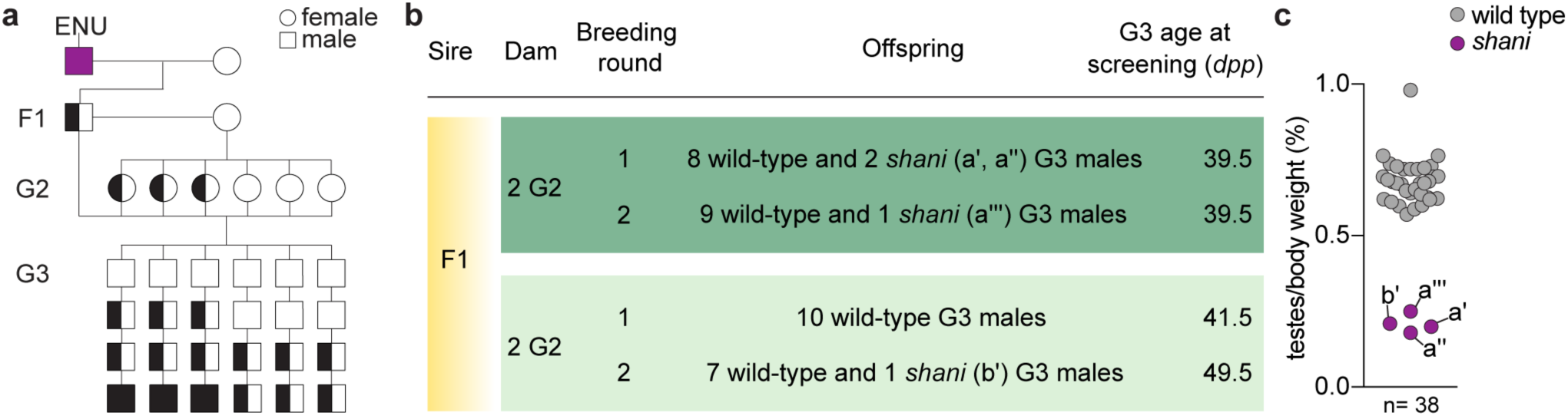
Mice from the ENU-induced mutant line *shani* have meiotic defects. **a** Breeding scheme. Male mice (C57BL/6J strain) were mutagenized and crossed to wild-type females (FVB/NJ strain) to produce founder (F1) males that were potential mutation carriers. Each F1 male was then crossed to wild-type females (FVB/NJ strain) to generate second generation (G2) offspring and G2 daughters were crossed back to their F1 sire to generate third-generation (G3) offspring. G3 males were screened for meiotic defects. If an F1 male was a carrier of a single autosomal recessive mutation of interest, roughly half of his G2 daughters were expected to be carriers and roughly one eighth of G3 males were expected to be homozygous. Un-filled shapes represent animals that are wild-type for a mutation of interest, half-filled shapes are heterozygous carriers, and filled shapes are homozygotes. **b** Screen results for the *shani* line. The F1 male was harem-bred to four G2 females, yielding 38 G3 males that displayed either a wild-type or *shani* phenotype (a′, a′′, a′′′, b′). Due to the harem-breeding setup we were unable to determine the proportion of G3 animals birthed from any given G2 female. **c** Testes-to-body-weight ratios for G3 mice screened (listed in panel **b**). Total number of G3 mice screened (n) is indicated and *shani* mutants (a′, a′′, a′′′, b′) are annotated.

We screened third-generation (G3) male offspring for meiotic defects by measuring testes weights. In adult wild-type mice, normal spermatogenesis results in testes populated with spermatogenic cell types spanning all stages of spermatocyte development. Mice with meiotic defects, such as failure to initiate or complete meiotic recombination, display spermatogenic arrest with apoptotic elimination of spermatocytes, resulting in a reduction in testes size or hypogonadism (de Rooij and de Boer 2003). To control for low testes weight due to poor growth and reduction in overall body size, we also measured the body weight and analyzed data as testes-to-body-weight ratios.

In an F1 founder line we named *shani*, four of the G3 males screened (**Fig. 1b**) showed a 69.6% reduction in testes-to-body-weight ratio compared to littermates (mean ratios were 0.21% for *shani* mutants and 0.69% for wild-type and heterozygous animals; p<0.01, one-sided t-test; **Fig. 1c**).

### shani maps to a missense mutation in the Spata22 gene

To identify the causative mutation, we generated additional mutant animals and performed whole-exome sequencing of DNA from five G3 *shani* mutants. Then we searched for homozygous variants that were shared between all five mutants (**Fig. 2a**). This revealed three un-annotated DNA sequence variants (**Fig. 2a**). Two variants were located within introns lacking sequencing read coverage in ENCODE long RNA-sequencing data from adult wild-type testis, making these unlikely phenotype-causing mutations. The third was an exonic variant located in the predicted *Spata22* coding sequence and was spanned by testis transcripts (**Fig. 2a, b**).

**Fig. 2.**
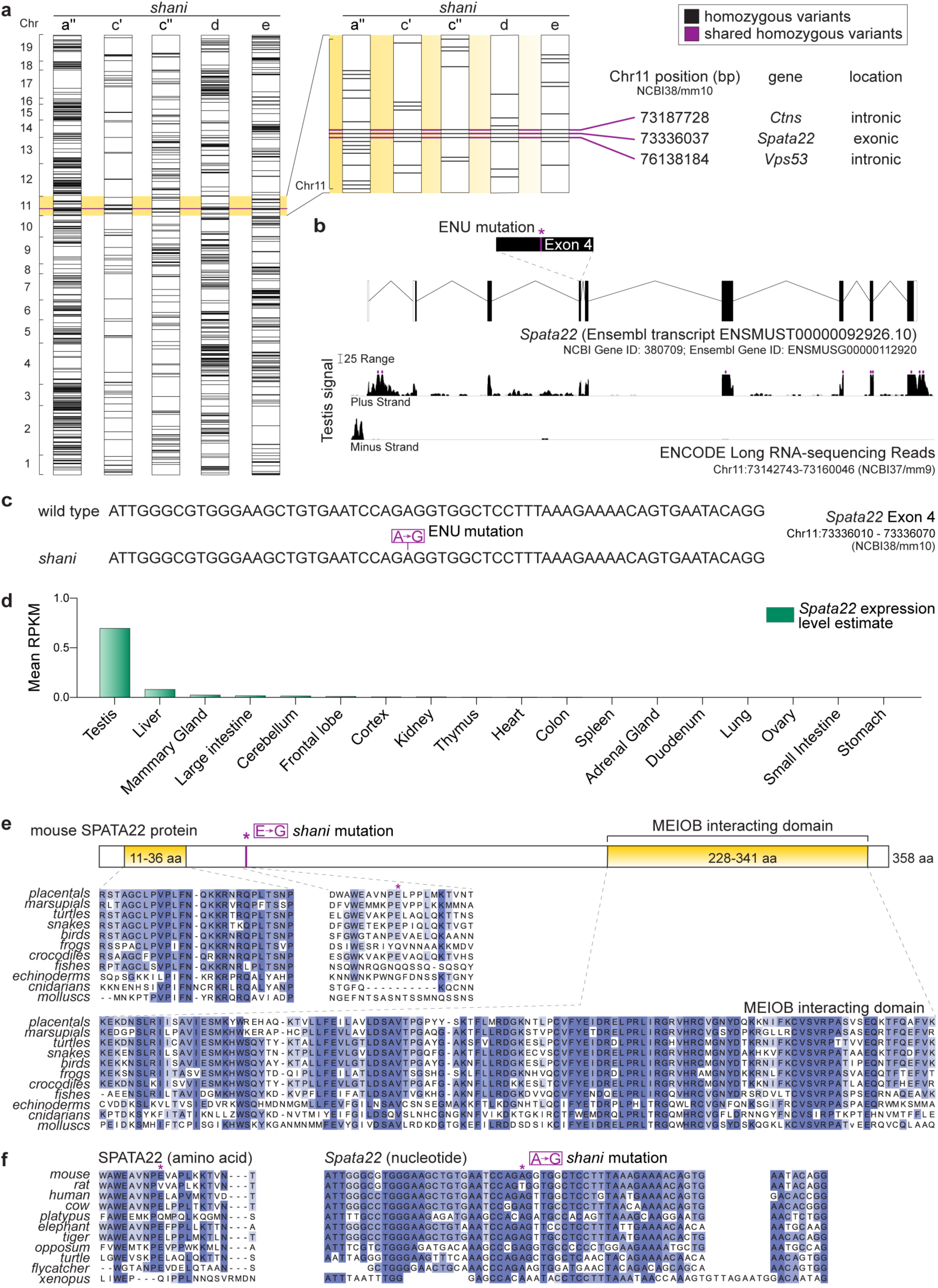
*shani* mice harbor a point mutation in *Spata22*. **a** Autosomal homozygous variants in five G3 *shani* mutants (a′′, c′, c′′, d, e) identified by whole-exome sequencing. Variants depicted exclude known between-strain SNPs. Vertical axis reflects variant index and chromosomes are not to scale. A detailed view of Chr11 variants is shown on the right. Three homozygous variants that are shared between mutants are highlighted in fuchsia with positions, gene names and genic locations listed. **b** Top: Schematic of *Spata22* (as predicted by Ensembl release 94) with protein-coding regions depicted as filled boxes. Location of the ENU-induced lesion is shown. Bottom: The density of ENCODE long RNA-sequencing reads (release 3) from adult testis within a window spanning from 500 bp upstream to 500 bp downstream of *Spata22*. The vertical viewing range is 0–50; read densities exceeding this range are overlined in fuchsia. **c** The *shani* allele of *Spata22*. **d** *Spata22* transcript expression level estimate (mean of reads per kilobase per million mapped reads (RPKM) values provided by ENCODE) in adult tissues. **e** Top: Schematic of mouse SPATA22 showing locations of conserved domains (yellow boxes) and the *shani* mutation. Bottom: Clustal Omega alignment of SPATA22 consensus amino acid sequences. Each line is a consensus sequence for the group and was derived using at least four species per group. Groups used are: placentals (Eutheria), marsupials (Metatheria), turtles (Testudines), snakes (Serpentes), birds (Aves), frogs (Anura), crocodiles (Crocodilia), fishes (Actinopterygii), echinoderms (Echinodermata), cnidarians (Cnidaria), molluscs (Mollusca). **f** Clustal Omega alignment of SPATA22 amino acid and *Spata22* nucleotide sequences from tetrapods corresponding to exon 4 of mouse *Spata22*. Location of the *shani* mutation is indicated. Species used are: mouse (*Mus musculus*), rat (*Rattus norvegicus*), human (*Homo sapiens*), cow (*Bos taurus*), platypus (*Ornithorhynchus anatinus)*, elephant (*Loxodonta Africana*), tiger (*Panthera tigris altaica*), opossum (*Monodelphis domestica*), turtle (*Pelodiscus sinensis*), flycatcher (*Ficedula albicollis*), xenopus (*Xenopus tropicalis*). Amino acid and nucleotide residues are shaded to reflect degree of conservation.

The *Spata22* variant is an A to G nucleotide transition at position Chr11:73336037, resulting in a missense mutation (glutamic acid to glycine) in exon 4 (**Fig. 2b, c**). *Spata22* RNA is expressed predominantly in the adult testis (**Fig. 2d**) and *Spata22*-null mice display meiotic defects with hypogonadism (Hays et al. 2017). This led us to surmise that this ENU-induced point mutation disrupts *Spata22* function and is the cause of the *shani* mutant phenotype.

To characterize the *shani* mutation, we examined conservation of *Spata22*. The mouse *Spata22* transcript encodes a protein of 358 amino acids and 40.17 kDa (presented schematically in **Fig. 2e**). Searches in protein databases revealed that placental mammals possess a similar SPATA22 protein: 358 to 364 amino acids in length with two well-conserved domains, a short motif located near the N-terminus and a longer region located near the C-terminus (yellow boxes in **Fig. 2e**). These domains were present in SPATA22 homologs from all vertebrates examined and were separated by a less conserved region of ∼190 amino acids. Using these domains to search, SPATA22 homologs were retrieved in invertebrates such as echinoderms, cnidarians and molluscs (**Fig. 2e, Table S1**). In several distant animals, e.g. arthropods (Arthropoda) and sponges (Porifera), a likely homolog restricted to the C-terminal domain could be retrieved but with substantial divergence from other homologs (**Table S1**). The conserved C-terminal domain corresponds to the region in mouse that is required for interaction with MEIOB, the functional collaborator and direct binding partner of SPATA22 (Xu et al. 2017; Ribeiro et al. 2018), suggesting that the SPATA22-MEIOB interaction is likely a conserved property of SPATA22.

The *shani* mutation lies outside of the two widely conserved SPATA22 protein domains (**Fig. 2e**). The mutation-containing region is poorly conserved in vertebrates (**Fig. 2e**), but shows higher conservation among tetrapods, particularly within mammals (**Fig. 2f**). We observed that this region is also well conserved at the nucleotide sequence level (**Fig. 2f**). Moreover, the mutated adenine is invariant in most mammalian genomes examined (except in rat), as well as in sauropsids and amphibians (**Fig. 2f**).

### Spata22^shani/shani^ mutants are sterile due to gametogenic failure

*Spata22*-null animals display a meiotic recombination defect, leading to meiotic prophase arrest and infertility (Hays et al. 2017), so we evaluated these phenotypes in mice carrying the *Spata22*^*shani*^ allele. Heterozygotes had normal fertility and transmitted the *shani* mutation in a Mendelian ratio (21.4% *Spata22*^*+/+*^, 49.7% *Spata22*^*shani/+*^ and 28.7% *Spata22*^*shani/shani*^ from heterozygote × heterozygote crosses; n = 219 mice; p=0.31, Chi-squared test). *Spata22*^*shani/shani*^ animals did not display obvious somatic defects. However, homozygous mutant males and females were sterile: none of the three *Spata22*^*shani/shani*^ males tested sired progeny when bred with wild-type females for 13**–**17 weeks and no pregnancies were observed from crosses of three homozygous mutant females to wild-type males (bred for 17 weeks).

In histological sections of adult testes, seminiferous tubules from control animals had the full array of spermatogenic cells, including spermatocytes and spermatids, as expected (**Fig. 3a**). In contrast, tubules from *Spata22*^*shani/shani*^ were greatly reduced in diameter and contained only early spermatogenic cells, with no post-meiotic germ cells visible (**Fig. 3a**). Mutants contained abnormal tubule types: some tubules (‘Em’ in **Fig. 3a**) had only a single layer of Sertoli cells and presumptive spermatogonia along the tubule perimeter; and some also had spermatogenic cells that appeared apoptotic (‘Apo’ in **Fig. 3a**). Increased apoptosis was confirmed by elevated TUNEL staining in mutants (**Fig. 3b, c**). These patterns are consistent with spermatocyte arrest and apoptosis during pachytene stage of meiotic prophase, occurring possibly at stage IV of the seminiferous epithelial cycle, which is typical of mutants with meiotic recombination defects (for example *Dmc1*^*-/-*^ (Barchi et al. 2005)) and is also observed in *Spata22*-null males (Hays et al. 2017).

**Fig. 3.**
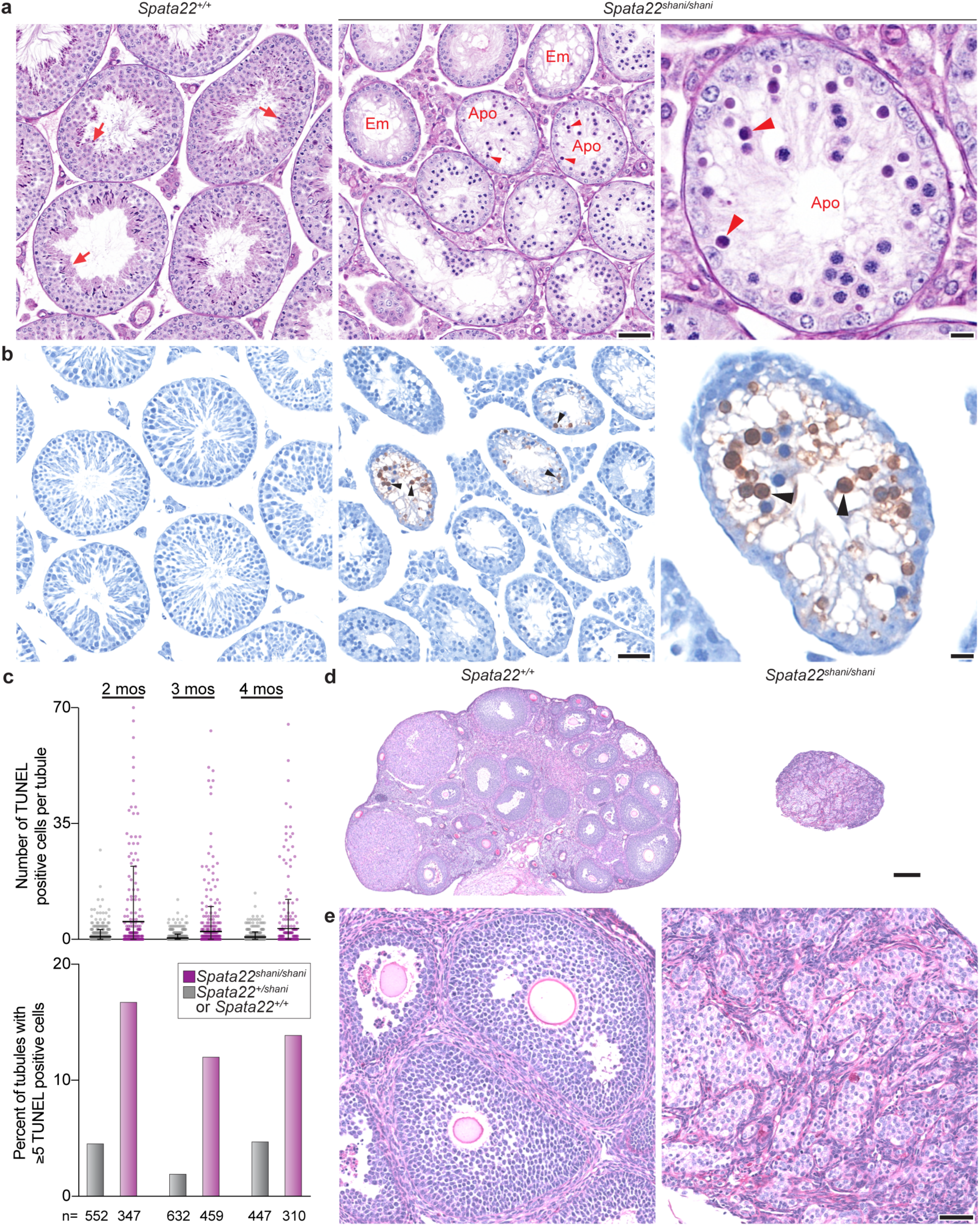
shani allele of Spata22 leads to gametogenic failure. **a** Periodic acid-Schiff (PAS)-stained sections of Bouin’s-fixed testes from a 3-month-old *Spata22*^*shani/shani*^ male and a wild-type littermate. Arrows indicate post-meiotic germ cells (spermatids) and arrowheads point to spermatocytes with an apoptotic morphology (condensed and/or fragmented). In the mutant, examples are indicated of apoptotic (“Apo”) tubules containing cells with apoptotic morphology and emptier-looking (“Em”) tubules harboring only cells with spermatogonia-like morphology and Sertoli cells. **b** TUNEL assay on testis sections from mice shown in panel **a**. Black arrowheads point to TUNEL-positive cells (stained dark brown). **c** Quantification of TUNEL assays performed on testis sections of *Spata22*^*shani/shani*^ mutants aged 2–4 months (mos) and their wild-type or heterozygous littermates. The number of seminiferous tubules counted from each animal is indicated (n) and lines in the dot plot depict mean ± SD. **d** and **e** Paraformaldehyde (PFA)-fixed, PAS-stained ovary sections from a 2-month-old *Spata22*^*shani/shani*^ female and a wild-type littermate. In panels **a** and **b**, the scale bars represent 50 µm and 10 µm in the lower and higher magnification images, respectively. In panels **d** and **e**, they represent 200 µm and 50 µm, respectively.

Defects in meiotic recombination in females cause elevated oocyte loss (Hunter 2017) and *Spata22*-null adult females have shrunken ovaries lacking oocytes (Hays et al. 2017). Meiotic recombination in females initiates during fetal development and arrests soon after birth, at which time oocytes mature into follicles that further develop during the reproductive life of the female (Peters 1969; Hunter 2017). As expected, adult wild-type ovaries had abundant developing follicles. In contrast, ovaries from adult *Spata22*^*shani/shani*^ females were dramatically smaller compared to wild-type littermates, and no developing follicles were visible (**Fig. 3d, e**), like in *Spata22*-null and consistent with the observed infertility.

### Spata22^shani/shani^ spermatocytes are defective in meiotic DSB repair and chromosome synapsis

Our initial findings showed that *Spata22*^*shani/shani*^ have spermatocyte arrest, apoptosis and infertility phenotypes similar to that observed in *Spata22*-null animals (Hays et al. 2017). We next examined meiotic recombination events, specifically DSB formation, repair and chromosome synapsis. We immunostained chromosome spreads of spermatocytes for SYCP3, a component of the chromosome axes (Lammers et al. 1994; Zickler and Kleckner 2015), and for γH2AX, a phosphorylated form of histone variant H2AX that is generated in response to DSBs (Mahadevaiah et al. 2001) (**Fig. 4a**). In normal meiosis, sister chromatids begin to form a SYCP3-containing protein axis from which chromatin loops emanate during the leptotene stage of meiotic prophase. During the zygotene stage, axes elongate and homologous chromosome axes begin to align and form stretches of the synaptonemal complex (SC), a tripartite structure comprising the homologous axial elements and the central region connecting them. The SC elongates and connects homologous chromosomes along their lengths in the pachytene stage. The SC disassembles during the diplotene stage (**Fig. 4a**). DSB formation occurs primarily during leptonema and zygonema, yielding nucleus-wide γH2AX staining. The γH2AX staining disappears progressively as recombination proceeds and is largely gone from autosomes during pachynema. γH2AX is also present on the X and Y chromosomes in the sex body during pachynema and diplonema (**Fig. 4a**).

**Fig. 4.**
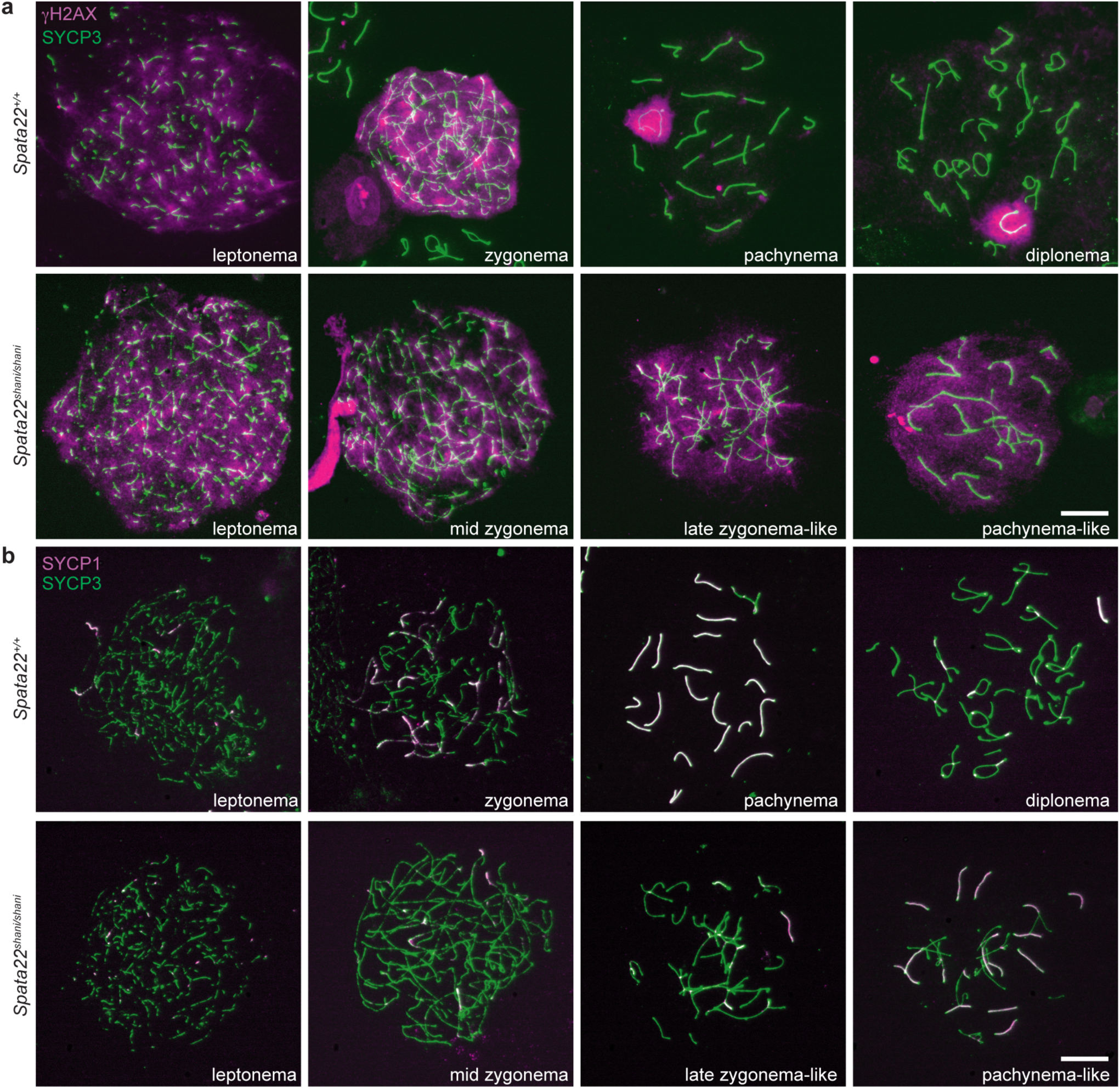
Meiotic prophase progression is impaired in *Spata22^shani/shani^*. Representative chromosome spreads from wild-type and *Spata22*^*shani/shani*^ spermatocyte nuclei assessing γH2AX (**a**) and SC formation (**b**). Prophase stages are indicated and mutant spermatocytes were classified based on SYCP3 patterns. Scale bars represent 10 µm.

*Spata22*^*shani/shani*^ spermatocytes with SYCP3 staining patterns indicative of leptonema and mid zygonema had pan-nuclear γH2AX staining, which is consistent with these cells having initiated meiotic recombination via formation of DSBs (**Fig. 4a**). *Spata22*^*shani/shani*^ spermatocytes with SYCP3 staining patterns indicative of later stages appeared abnormal, however. Cells with staining showing elongated axes that have begun to align (late zygonema-like) were abundant and retained pan-nuclear γH2AX staining (**Fig. 4a**). There were also rare cells where few homologous chromosomes appeared to be connected along their entire lengths (pachynema-like) (**Fig. 4a**).

We also tracked synapsis by co-immunostaining for SYCP3 and SYCP1, a SC central region protein (de Vries et al. 2005; Zickler and Kleckner 2015). In wild type, SYCP1 staining appears during zygonema as cells begin to build the SC, it marks autosomes along their lengths during pachynema when the SC connects homologous chromosomes and is largely absent from the autosomes in diplonema when the SC is disassembled (**Fig. 4b**). In *Spata22*^*shani/shani*^ littermates, leptonema and mid zygonema appeared normal. But later-stage spermatocytes (late zygonema-like and pachynema-like) had SYCP1 patterns indicative of synapsis defects: numerous *Spata22*^*shani/shani*^ spermatocytes possessed a mix of partially synapsed, fully synapsed and asynaptic chromosomes, along with chromosome tangles (i.e., a mix of synapsed and unsynapsed axes with partner switches indicative of nonhomologous synapsis; **Fig. 4b**). These patterns are typical of recombination- and synapsis-defective mutants such as *Dmc1*^*-/-*^ (Pittman et al. 1998; Yoshida et al. 1998), and are comparable to that observed in *Spata22*-null spermatocytes (Hays et al. 2017). We conclude that *Spata22*^*shani/shani*^ spermatocytes have a defect in meiotic DSB repair and chromosome synapsis, leading to meiotic prophase arrest and apoptosis. This culminates in the absence of postmeiotic cells, hypogonadism and sterility.

### Recombination dynamics in Spata22^shani/shani^ mutants

To evaluate the molecular characteristics of the *Spata22*^*shani/shani*^ cells with a meiotic recombination defect, we examined behaviors of recombination proteins by immunostaining spread spermatocyte chromosomes. SYCP3 staining patterns were used to define meiotic prophase stages and only recombination protein foci overlapping with SYCP3-positive axes were counted. We first analyzed DMC1 strand-exchange protein that assembles on ssDNA formed at DSBs and is required for homology search and strand invasion during meiotic recombination (Brown and Bishop 2014). In normal meiosis, DMC1 foci appear in leptonema, accumulate to maximal levels in early to mid zygonema, then decline as DSB repair proceeds through pachynema (**Fig. 5a, b**). In *Spata22*^*shani/shani*^, DMC1 foci accumulate during early stages but prematurely decline in mid zygonema and are severely reduced in late zygonema compared to wild type (**Fig. 5a, b**; mean foci count of mid zygonema and late zygonema-like *Spata22*^*shani/shani*^ = 96 and 27, respectively; mean foci count of mid zygonema and late zygonema wild type = 154 and 101, respectively). These findings are consistent with previous observations of DMC1 behavior in *Spata22*-mutant rats as well as in mice lacking MEIOB, the functional collaborator and direct binding partner of SPATA22 (Souquet et al. 2013; Ishishita et al. 2014). *Spata22*^*shani/shani*^ DMC1 foci counts in late leptonema and early zygonema, although not statistically significant, were lower compared to wild type (**Fig. 5b**), suggesting that DMC1 accumulation may also be modestly affected at these stages.

**Fig. 5.**
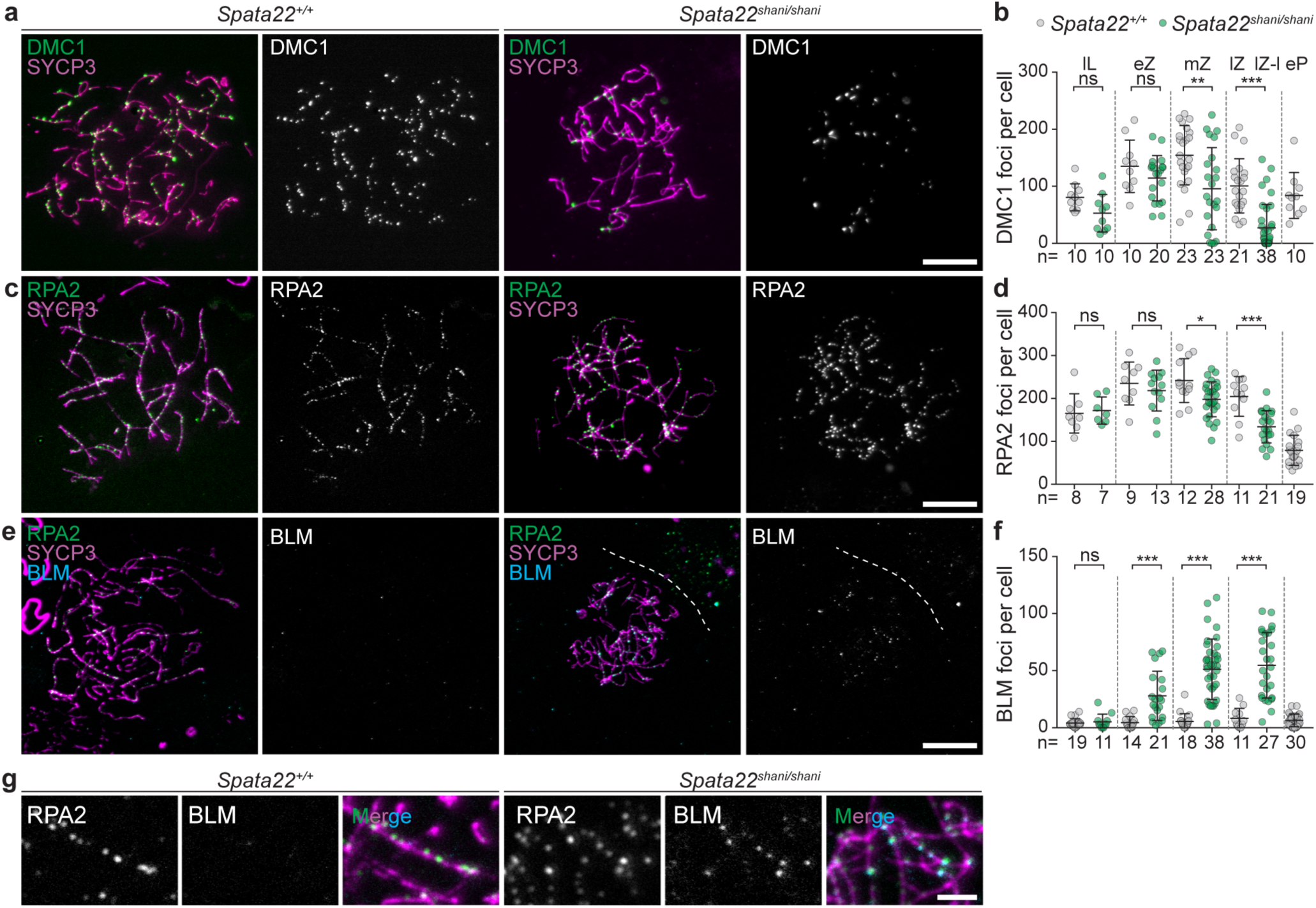
*Spata22^shani/shani^* spermatocytes show precocious DMC1 loss and abnormal BLM accumulation. **a, c, e** Immunostained chromosome spreads from wild-type and *Spata22*^*shani/shani*^ spermatocyte nuclei. Images of representative zygotene stages are presented. **b, d, f** Quantification of DMC1, RPA2 and BLM foci. Each point is the count from one spermatocyte and number of spermatocytes analyzed is indicated (n). Lines depict mean ± SD and the results of two-tailed Mann-Whitney tests are shown: ns is not significant (p>0.05); *p<0.05, **p<0.01, ***p<0.001. Meiotic prophase stages are as follows: late leptonema (lL), early zygonema (eZ), mid zygonema (mZ), late zygonema (lZ), early pachynema (eP) and late zygonema-like (lZ-l) in mutant. Spermatocytes were classified based on SYCP3 patterns. **g** Higher magnification view of immunostaining shown in **e**. Scale bars represent 10 µm in panels **a, c, e** and 2 µm in panel **g**.

We also examined RPA2, which is a subunit of the ubiquitous, trimeric ssDNA-binding complex RPA (Wold 1997). RPA is present on ssDNA-containing intermediates during DSB metabolism. In wild type, RPA2 foci form during leptonema, increase in early and mid zygonema, and then decrease as DSB repair progresses in late zygonema and early pachynema (**Fig. 5c, d**). *Spata22*^*shani/shani*^ cells accumulate RPA2 foci during early stages and arrest during zygonema with RPA2 foci present (**Fig. 5c, d**). This is as observed in *Spata22*-mutant rats and *Meiob*^*-/-*^ mice (Luo et al. 2013; Souquet et al. 2013; Ishishita et al. 2014), suggesting that arrested cells likely possess ssDNA-containing intermediates. We observed that RPA2 foci counts in mid zygonema and late zygonema were reduced compared to wild type (**Fig. 5d**; mean foci count of mid zygonema and late zygonema-like *Spata22*^*shani/shani*^ = 198 and 134, respectively; mean foci count of mid zygonema and late zygonema wild type = 242 and 205, respectively), likely due to defects in recombination intermediate processing during these stages.

We further investigated localization of the RecQ helicase family protein, Bloom’s syndrome mutated (BLM). BLM and its orthologs process multiple types of recombination intermediates (Singh et al. 2009; De Muyt et al. 2012). BLM has been previously reported to form discrete foci on chromosome axes during wild-type meiosis (Moens et al. 2000; Holloway et al. 2010) and to accumulate in recombination-deficient mutants (Holloway et al. 2008; Holloway et al. 2010). In our hands, BLM was barely detectable in most wild-type spermatocytes, with few cells forming clear axis-associated foci that met our immunofluorescence cutoff (**Fig. 5e, f**). In contrast to wild type, *Spata22*^*shani/shani*^ had bright and numerous BLM foci on chromosome axes (**Fig. 5e, f**). RPA has been shown to stimulate BLM activity *in vitro* and to colocalize with BLM during meiosis (Walpita et al. 1999; Brosh et al. 2000), therefore we asked if the BLM foci we see in mutants colocalized with RPA. Most BLM foci also had an RPA2 focus present (**Fig. 5g**). BLM foci were first detected in early zygonema and levels continued to rise as cells progressed to mid zygonema and late zygonema-like stages (mean foci count of late zygonema-like *Spata22*^*shani/shani*^ = 55). Our results suggest that in *Spata22*^*shani/shani*^, either there is increased BLM recruitment (for example, an increase in the presence or accessibility of BLM targets) or that BLM is abnormally retained. Taken together, our results indicate a failure to repair DSBs appropriately and/or the presence of aberrant DNA structures in *Spata22shani/shani*.

### SPATA22 protein is not detected in Spata22^shani/shani^ testes

During normal meiosis, SPATA22 appears on chromosome axes as foci that match DSB dynamics: SPATA22 foci are first detectable during late leptonema, focus numbers increase progressively through zygonema, then decrease through pachynema and are no longer detected during diplonema (**Fig. 6a, b**)(Ishishita et al. 2014; Hays et al. 2017). SPATA22 patterns are similar to those for MEIOB, as expected for members of a protein complex (Luo et al. 2013; Souquet et al. 2013).

**Fig. 6.**
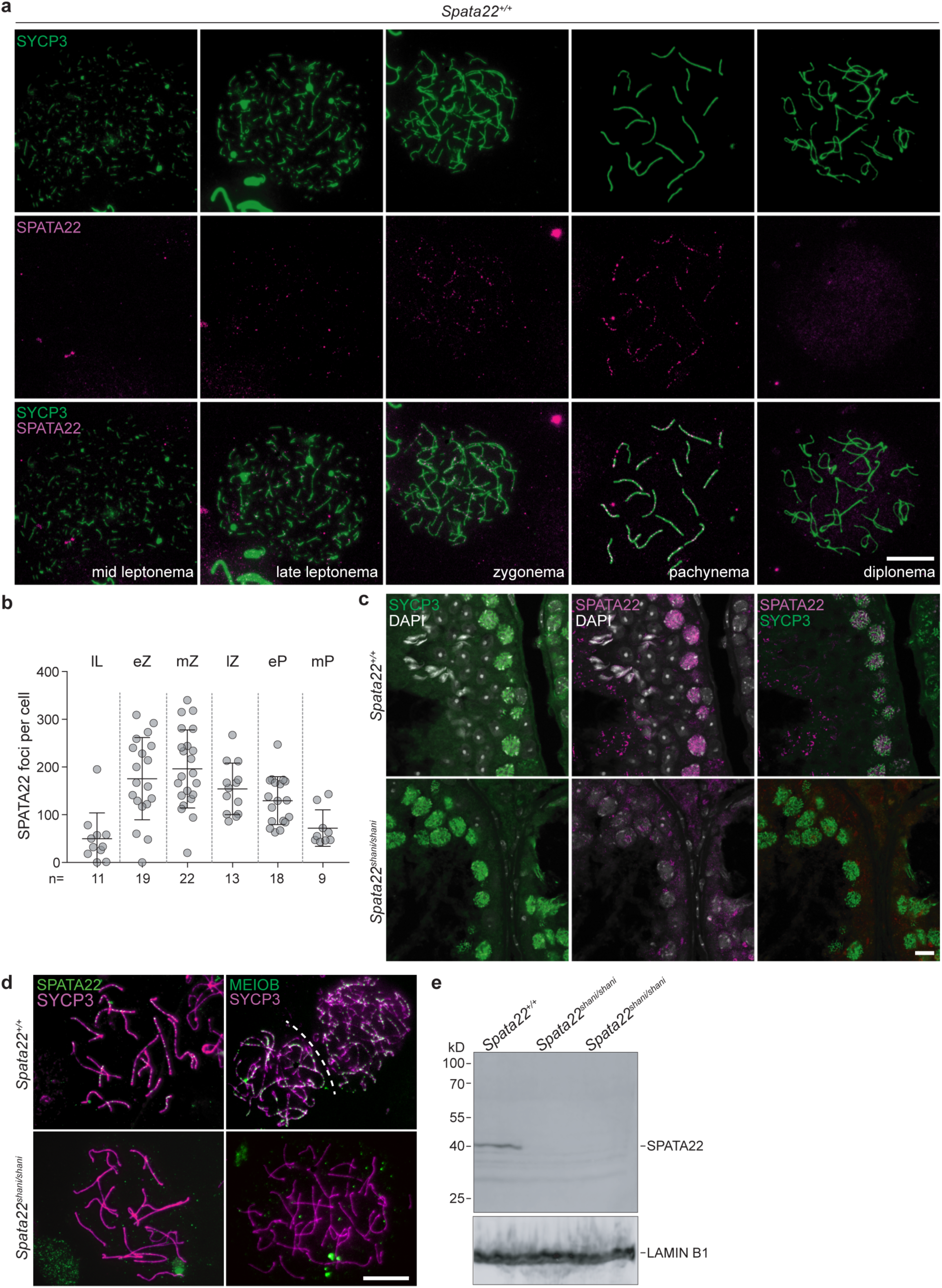
SPATA22 localization and protein levels in wild-type and *Spata22^shani/shani^* testes. **a** Representative wild-type chromosome spreads from spermatocyte nuclei. **b** Quantification of SPATA22 foci in wild type. Each point is the count from one spermatocyte and number of spermatocytes analyzed is indicated (n). Lines depict mean ± SD. Meiotic prophase stages are abbreviated as in **Fig. 5b**; mid pachynema (mP). **c** SYCP3 and SPATA22 immunofluorescence on testis sections from 11-month-old *Spata22*^*shani/shani*^ and wild-type males. Sections containing cells with similar SYCP3 profiles are shown and DNA is stained with DAPI. **d** Defective SPATA22 and MEIOB localization in *Spata22*^*shani/shani*^. Representative chromosome spreads with similar SYCP3 profiles are shown (late zygonema for *Spata22*^*+/+*^ and cells arrested at late zygonema-like for *Spata22*^*shani/shani*^). **e** Western blot analysis of SPATA22 protein levels in whole-testis lysates from two 16-dpp *Spata22*^*shani/shani*^ and a wild-type littermate. Scale bars represent 10 µm in panels **a, c, d**.

Since the meiotic recombination defect and spermatocyte arrest in *Spata22*^*shani/shani*^ phenocopied a null mutation, we asked whether SPATA22 protein was properly localized. In testis sections, SPATA22 staining was prominent in SYCP3-positive spermatocytes in wild type, but little to no staining was observed in *Spata22*^*shani/shani*^ littermates (**Fig. 6c**). On spread spermatocyte chromosomes, whereas SPATA22 foci were visible along axes in wild type, similarly staged cells from *Spata22*^*shani/shani*^ littermates showed only weak background staining that did not overlap with axes (**Fig. 6d**, left panels).

To discern whether the lack of staining in *Spata22*^*shani/shani*^ spermatocytes was due to protein mis-localization or lack of protein, we performed western blot analysis on whole testis lysates. Because mutants undergo spermatocyte arrest, we examined juvenile mice aged 16 dpp. At this age spermatocytes in meiotic prophase are abundant (Bellve et al. 1977), however we expected that programmed cell death had not yet significantly altered the relative sizes of germ cell subpopulations. While SPATA22 could be detected in wild type at the expected size, no bands were detected in the mutant using a polyclonal antibody (**Fig. 6e**). Thus it is likely that the *shani* mutation leads to lack of protein production or to protein destabilization.

Absence of SPATA22 strongly impairs the stability of its partner, MEIOB (Luo et al. 2013; Hays et al. 2017). Consequently, MEIOB is not detected on chromosome axes in spermatocytes from SPATA22-null mice (Luo et al. 2013; Hays et al. 2017). Similarly, we could not detect MEIOB on chromosome axes in *Spata22*^*shani/shani*^ spermatocytes (**Fig. 6d**, right panels). We conclude that the meiotic recombination defect in *Spata22*^*shani/shani*^ spermatocytes is a consequence of decreased SPATA22 protein levels and decreased SPATA22-MEIOB complex on chromosome axes.

### The shani mutation leads to defective Spata22 splicing and RNA instability

To determine how the *shani* mutation leads to decreased SPATA22 protein levels, we examined *Spata22* mRNA by reverse transcription-PCR (RT-PCR) using primers directed to the 5’ and 3’ UTRs (primer set X; **Fig. 7a, b**). *Spata22*^*+/+*^, *Meiob*^*+/+*^ and *Meiob*^*+/-*^ adult testes yielded a 1239-bp band, consistent with the predicted length of wild-type *Spata22* mRNA. PCR products from *Spata22*^*shani/shani*^ testes were shorter, however (**Fig. 7b**).

**Fig. 7.**
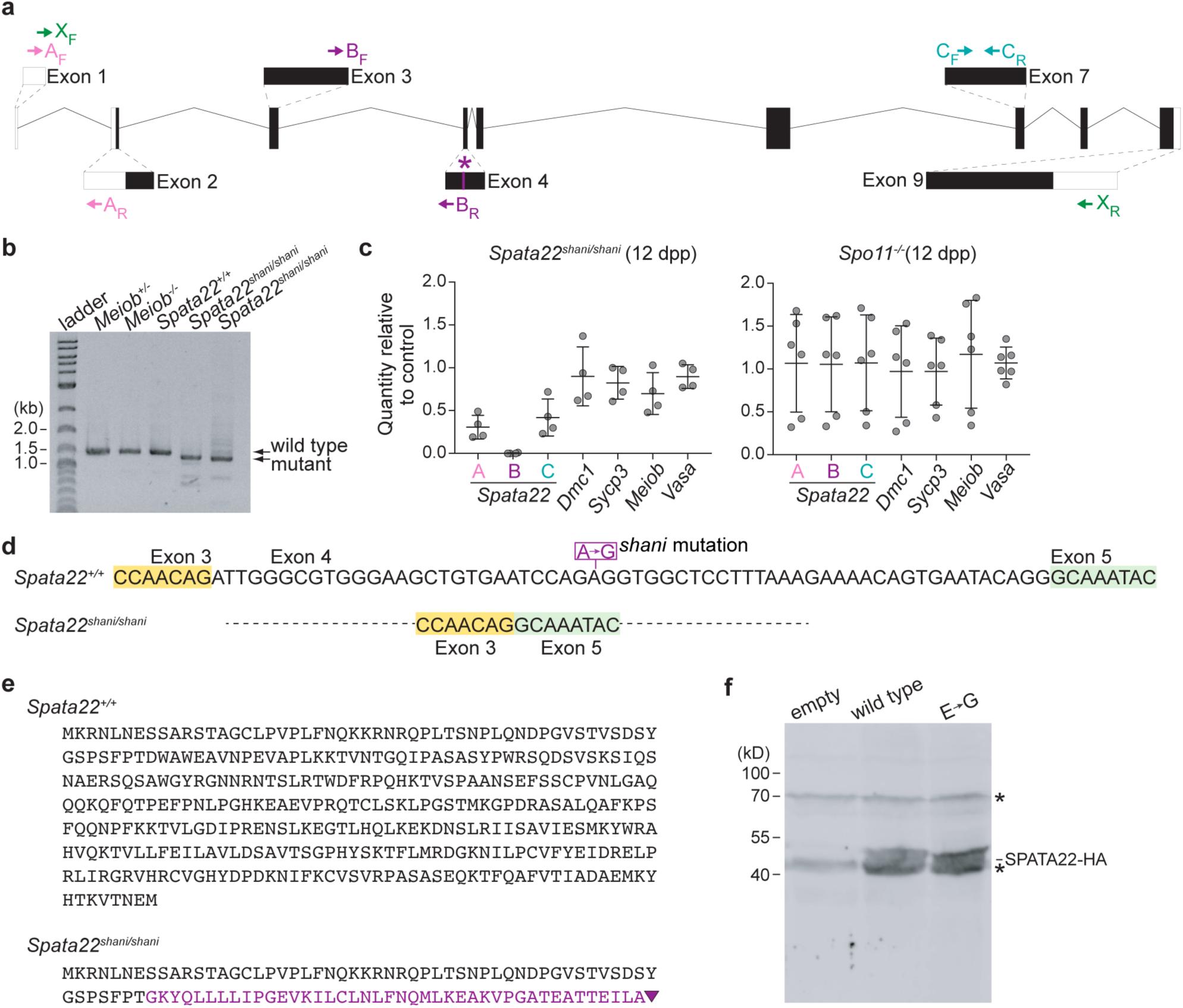
The *shani* mutation causes defective splicing of *Spata22* RNA. **a** Schematic of the wild-type *Spata22* transcript (same as in **Fig. 2b**) showing locations of the *shani* mutation (asterisk) and RT-PCR primers. Boxes represent exons with coding regions shown as filled boxes and lines represent introns. Primers X_F_ (green; spanning exon-exon junction between exons 1 and 2) and X_R_ (green; located within exon 9) flank the *Spata22* coding region, primers A_F_ and A_R_ (pink; located within exons 1 and 2, respectively) flank the exon-exon junction between exons 1 and 2, primers B_F_ (fuchsia; located within exon 3) and B_R_ (fuchsia; spanning exon-exon junction between exons 3 and 4) amplify the exon-exon junction region between exons 3 and 4, primers C_F_ and C_R_ (blue) are located within exon 7. **b** RT-PCR analysis of *Spata22* in whole-testis RNA samples extracted from 3-month-old *Spata22*^*shani/shani*^ males, a 15-month-old *Meiob*^*-/-*^ male and wild-type or heterozygous littermates. PCR amplification was done using primers flanking the *Spata22* coding region (primers X; shown in green in **a**) and analyzed by gel electrophoresis. Black arrows indicate amplified products. **c** Quantitative RT-PCR analysis of whole-testis RNA samples from 12-dpp *Spata22*^*shani/shani*^ and 12-dpp *Spo11*^*-/-*^ mice. Data points represent individual mice normalized to wild-type or heterozygous littermates. Lines depict mean ± SD. Transcripts analyzed are noted along the x-axis. Three regions of *Spata22* (shown in **a**) were analyzed: exon 1-exon 2 junction (A), exon 3-exon 4 junction (B) and internal region of exon 7 (C). **d** Nucleotide sequence of *Spata22* mRNA spanning exon 4 and flanking regions in wild type and *Spata22*^*shani/shani*^. *Spata22* coding region was amplified as described in **b**, gel purified and products were analyzed by Sanger sequencing. Location of the *shani* mutation is indicated. **e** Predicted amino acid sequence of SPATA22 in wild type and *Spata22*^*shani/shani*^. Wild-type amino acid sequence is shown in black. Exon 4 loss in mutants is predicted to result in a frameshift, alternative amino acid sequence (fuchsia) and a premature stop codon (triangle). **f** Western blot analysis of HA-tagged human SPATA22 constructs expressed in HEK-293 cells. Constructs carrying wild-type *Spata22* coding sequence, coding sequence with the *shani* mutation (E→G), or an empty construct were expressed and blots probed with anti-HA antibody. Location of SPATA22-HA of wild-type length is indicated and asterisks denote non-specific bands.

We next assessed mRNA levels by quantitative RT-PCR using three primer sets spanning segments of *Spata22* (primer sets A, B, C; **Fig. 7a**). We examined animals at 12 dpp when *Spata22*-expressing early meiotic cells are abundant (Bellve et al. 1977) and spermatogenic arrest is not expected to have greatly affected the relative population sizes of germ cell subtypes. We also examined 12 dpp-old *Spo11*^*-/-*^ mice that lack DSB formation and show a similar arrest as *Spata22*^*shani/shani*^ (Baudat et al. 2000; Romanienko and Camerini-Otero 2000).

*Spata22* mRNA regions A and C had reduced amplification in 12 dpp-old *Spata22*^*shani/shani*^ (relative quantities were 0.31 and 0.42, respectively; **Fig. 7c**) while control meiotic transcripts were normal (*Dmc1, Sycp3, Meiob* and *Vasa*; **Fig. 7c**), suggesting that *Spata22* mRNA level is decreased in *Spata22*^*shani/shani*^. All transcripts assayed were normal in 12 dpp-old *Spo11*^*-/-*^ mice, confirming that the reduced amplification of *Spata22* mRNA is not due to spermatogenic arrest. Intriguingly, *Spata22* mRNA region B did not yield an amplification product in 12 dpp-old *Spata22*^*shani/shani*^ (**Fig. 7c**).

We performed Sanger sequencing on the wild-type *Spata22* cDNA and shorter variant produced in *Spata22*^*shani/shani*^ (**Fig. 7b**). cDNA sequence from wild type was as expected but cDNA from mutant testes completely lacked exon 4, with exon 3 and exon 5 spliced together to create a new exon-exon junction (**Fig. 7d**). This is consistent with the lack of amplification of region B (exon 3-exon 4 junction; **Fig. 7a, c**). Together these results suggest that the *shani* mutation leads to expression of a less stable and shorter variant of *Spata22* mRNA that lacks exon 4.

Loss of exon 4 predicts a frameshift, resulting in a stop codon within exon 5 and the formation of a truncated protein of 99 amino acids and ∼11 kDa, with the N-terminal 57 amino acids of wild type sequence (black in **Fig. 7e**) and the remaining amino acids of alternative sequence (fuschia in **Fig. 7e**). We were unable to detect a truncated SPATA22 protein by western blot or immunolocalisation in *Spata22*^*shani/shani*^, but we do not know whether our SPATA22 antibody would recognize the predicted truncation.

The *shani* mutation on exon 4, if translated, is predicted to lead to a glutamic acid to glycine substitution. We wondered whether the *shani* mutation would compromise protein stability if exon 4 were to be retained on the *Spata22* transcript. To test this, we transfected human cells with either a plasmid expressing wild-type *Spata22* cDNA, or a plasmid expressing *Spata22* cDNA that harbors the *shani* mutation, or an empty plasmid as a control. Western blot analysis of SPATA22 showed a ∼40 kDa band in wild type, as expected (**Fig. 7f**). Cells expressing *Spata22* cDNA with the *shani* mutation also had a ∼40 kDa band which was similar in intensity to that in wild type (**Fig. 7f**), indicating that the *shani* mutation, when present on a transcript that does not need to be spliced, does not affect production or stability of the SPATA22 protein.

## Discussion

This study describes a novel point-mutated allele of the meiotic recombination gene *Spata22* using a phenotype-based forward-genetics screen. *Spata22*^*shani/shani*^ mutant spermatocytes appear to initiate DSB formation normally during meiotic prophase, but show defects in DSB repair and synapsis, leading to spermatocyte arrest, apoptosis, shrunken gonads and infertility. We performed molecular characterization of the *shani* mutation and show that it leads to defective splicing of the *Spata22* RNA, ultimately resulting in loss of detectable SPATA22 protein. Consistent with this, *Spata22*^*shani*^ and *Spata22*-null alleles have similar meiotic phenotypes (Hays et al. 2017). Our work illustrates the power of forward-genetic approaches in exploring the regulation of mammalian meiosis. Moreover, we describe a unique screen design and demonstrate its feasibility: we exploited a hallmark of meiotic recombination defects, hypogonadism, to design an efficient and simple readout based on testes weights. Also, we utilized a new mutation identification strategy that leverages recent genomic advances.

We show that SPATA22 has two well-conserved domains: an N-terminal segment and a longer region located near the C-terminus. Similar findings have been previously reported, although the precise boundary of the conserved C-terminal domain differs (La Salle et al. 2012). The N-terminus of SPATA22 has no predicted structure or known function. The conserved C-terminal region we describe here (228–341 amino acids) closely matches its predicted OB fold-containing domain (277–356 amino acids), which is essential for its interaction with MEIOB (Xu et al. 2017; Ribeiro et al. 2018). Like SPATA22, MEIOB is predicted to have OB fold-containing domains (Luo et al. 2013; Souquet et al. 2013; Ribeiro et al. 2016) and domain architectures of both proteins resemble those of RPA subunits (Luo et al. 2013; Ribeiro et al. 2016; Ribeiro et al. 2018). Also, both proteins interact and colocalize with RPA subunits (Luo et al. 2013; Souquet et al. 2013; Ishishita et al. 2014; Xu et al. 2017; Ribeiro et al. 2018). These findings lead to the hypothesis that the SPATA22-MEIOB complex collaborates with RPA to regulate meiotic recombination. The precise stoichiometry of the complex and whether it includes RPA subunits or other proteins remains unclear, however. The biochemical role of the SPATA22-MEIOB complex is also unknown. A truncated form of MEIOB has been reported to have exonuclease activity *in vitro* (Luo et al. 2013) and recent studies show that SPATA22-MEIOB can organize RPA-coated ssDNA *in vitro* (Ribeiro et al. 2018). How these activities relate to its meiotic recombination function *in vivo* is unclear.

We show that the initial formation of DMC1 foci appears unaffected in *Spata22*^*shani/shani*^ and these foci decrease in number as prophase progresses in mice lacking SPATA22; this was also shown previously for DMC1 and RAD51 in mice lacking MEIOB and for RAD51 in rats lacking SPATA22 (La Salle et al. 2012; Luo et al. 2013; Souquet et al. 2013; Ishishita et al. 2014). We found that in the absence of SPATA22-MEIOB, DMC1 foci numbers decrease faster than in wild type (this manuscript and (Souquet et al. 2013)). This was also observed for RAD51 in mice lacking MEIOB and rats lacking SPATA22 (Souquet et al. 2013; Ishishita et al. 2014). One study did not report an earlier loss of strand-exchange proteins (Luo et al. 2013), but they analyzed juvenile mice where recombination foci dynamics have been reported to differ (Zelazowski et al. 2017). We found that RPA2 foci number also decrease earlier than in wild type. Since RPA is thought to decorate multiple recombination intermediates, one possibility is that there is a defect in the formation or stability of a subpopulation of RPA foci. The reduction in RPA2 foci number was not described previously in *Spata22* or *Meiob* mutants (Luo et al. 2013; Souquet et al. 2013; Ishishita et al. 2014); differences in strain background may explain this discrepancy. The disappearance of DMC1 and RPA during wild-type meiosis is an indicator of repair progression, thus a simple interpretation of our data is that some repair processes may occur faster or earlier in mutants, at least at some sites.

We found that while RPA2 foci number decrease over the course of meiotic prophase progression in *Spata22* mutants, cells arrest with RPA2 foci present, similar to MEIOB mutants and SPATA22-mutant rats (Luo et al. 2013; Souquet et al. 2013; Ishishita et al. 2014). The identity of these RPA foci is unknown and may represent a combination of different wild-type and mutant states. Mutants also show persistent γH2AX staining (La Salle et al. 2012; Luo et al. 2013; Souquet et al. 2013; Ishishita et al. 2014; Hays et al. 2017). These results indicate that not all sites are properly repaired, if any. It is possible that strand invasion occurs and DMC1 is lost, but that one or more subsequent steps such as D-loop stability or DNA synthesis or second end capture are misregulated. It is also possible that strand-exchange proteins are destabilized or that their ssDNA substrates are reduced, modified or become less accessible. All these scenarios may lead to the presence of structures that would be recognized by BLM and could explain the dramatic BLM accumulation we see in mutants. BLM plays multiple roles to eventually prevent formation of potentially toxic structures: BLM disrupts RAD51 nucleoprotein filaments or D-loops, stimulates primer extension and promotes Holliday junction dissolution (Wu et al. 2001; Bugreev et al. 2007). Regardless of the underlying role of BLM in mutants, its accumulation reinforces the requirement of SPATA22-MEIOB in proper processing of recombination intermediates.

Recently, SPATA22 has been shown to immunoprecipitate in testis extracts with MEILB2/HSF2BP (Zhang et al. 2019). MEILB2 interacts with BRCA2, which promotes RAD51/DMC1 nucleoprotein filament formation, and *Meilb2*^*-/-*^ meiotic cells fail to form RAD51/DMC1 foci (Brandsma et al. 2019; Zhang et al. 2019). The authors propose that SPATA22-MEIOB may be involved in MEILB2 and BRCA2 recruitment to DNA, which would then facilitate RAD51/DMC1 loading (Zhang et al. 2019). This model predicts that RAD51/DMC1 would not accumulate in SPATA22-MEIOB mutants. We observe only a modest reduction in DMC1 foci numbers during leptonema and early zygonema, but we do not know whether the DMC1 foci we detect at any of these stages are properly organized, stable or functional. Our results support the idea that RAD51/DMC1 filament assembly, stability and disassembly involves a complex regulatory network with cross-talk between multiple factors. The molecular determinants of these interactions remain to be elucidated.

We performed molecular characterization of the *shani* lesion and uncovered surprising regulation of *Spata22* gene expression. The *shani* mutation alters nucleotide sequences within exon 4 that are crucial for accurate splicing of *Spata22*. The splicing regulation we document here has been reported in mammalian genes, where long introns and weakly defined exon-intron boundaries are supported by both exonic and intronic cis-acting elements that either stimulate or repress splicing. One prevalent exonic element, called ESE, serves as a binding site for splicing factors called serine/arginine-rich (SR) proteins to promote exon definition (Blencowe 2000; Cartegni et al. 2002). There are several instances where disease-causing mutations that result in alternative splicing have been proposed to affect ESEs and mutations that abrogate an ESE often cause exon-skipping (Blencowe 2000; Cartegni et al. 2002), akin to that observed in *shani* mutants. We propose that the *shani* mutation possibly alters an ESE element within exon 4 in the *Spata22* transcript, leading to skipping of exon 4 by the spliceosome, making this, to our knowledge, the first description of an ESE element in a murine meiotic gene. While ESE elements lack a well-defined consensus sequence, combinatorial *in vivo* and *in vitro* approaches have led to the identification of 6- to 8-nucleotide long ESE sequence motifs and been used to develop computational ESE-prediction tools (Blencowe 2000; Cartegni et al. 2002). Indeed, the CCAGAGG and CAGAGGT overlapping sequences encompassing the affected adenine in *shani* mutants are recognized as putative ESE motifs by the ESEfinder ESE-prediction web tool (Cartegni et al. 2003).

While the amino acid affected by the *shani* mutation lies outside of the well-conserved SPATA22 protein domains, the affected DNA nucleotide (adenine) appears to be conserved in most tetrapods examined, as is the entirety of exon 4. This indicates that the splicing regulation we describe above may not be restricted to mouse, but rather may be a conserved feature of *Spata22* gene expression regulation. Similar regulation may be important for controlling other meiotic genes, especially given the extensive gene expression modulations that occur during meiosis and germ cell development. Our findings highlight how coding single-nucleotide polymorphisms can deregulate/alter splicing events and lead to phenotypic variability or gene inactivation during development. Moreover, our isolation and characterization of the *shani* allele of *Spata22* demonstrates the power of forward genetic approaches in uncovering unexpected gene regulation in mammals.

## Materials and Methods

### Generation of shani mutants

Details of the mutagenesis and breeding for screening purposes are described elsewhere (Jain et al. 2017) and were similar to previously described methods (Caspary 2010; Probst and Justice 2010). We screened G3 males for meiotic defects at ≥35 dpp, focusing on meiotic recombination. By this age the testes weight reduction was deemed to be clearly discernable for known meiotic recombination mutants (Yoshida et al. 1998). Thus G3 mutants displaying arrest and spermatocyte elimination similar to that seen in typical meiotic recombination mutants were expected to be identifiable. Testes pairs were dissected, weighed along with the body, frozen in liquid nitrogen and stored at −80°C until samples from ∼24 or more G3 males had been collected. Then testes-to-body-weight ratios were evaluated side-by-side for all animals from any given line.

Mice with an obvious reduction in testes-to-body-weight ratio compared to littermates were subjected to secondary screening by histological examination. One frozen testis from potential mutant mice, along with one frozen testis from a wild-type littermate were used for histology and the second testis was reserved for DNA extraction. Immunohistochemical TUNEL assay was performed and seminiferous tubules were evaluated for the presence of TUNEL staining, depletion of seminiferous tubules and absence of spermatids.

Genotyping of *shani* animals was done by PCR amplification using *shani* F1 and *shani* R primers, followed by Sanger sequencing using *shani* F2 primer (oligonucleotide primer sequences are provided in **Table S2**) or by digestion of the amplified product with MnlI (NEB) or BstNI (NEB). The *shani* mutation eliminates an MnlI restriction site that is present in the wild-type sequence and creates a novel BstNI restriction site.

### shani mutation identification and exome sequencing

Genome assembly coordinates are from GRCm38/mm10. We performed whole-exome sequencing on five G3 *shani* mutants (a′′ in **Fig. 1b**; c′, c′′ from a fourth F1xG2 harem-breeding cage that was set up after the initial screen cross; d and e from a fifth and sixth harem-breeding cage set up after the initial screen cross). Genomic DNA was extracted from testes or tail biopsies as described (Jain et al. 2017) and whole-exome sequencing was performed at the MSKCC Integrated Genomics Operation as described (Jain et al. 2018) to generate approximately 80 million 100 bp paired-end reads. Read alignment, variant calling and variant annotation were done as described (Jain et al. 2017).

We expected that lines generated from distinct mutagenized males and mutant lines with distinct phenotypes would likely not share phenotype-causing variants. Thus variants called from distinct mutant lines sequenced in parallel with the *shani* mutant line were filtered out to identify variants that are private to the *shani* line (i.e., found in *shani* mutant mice but not in mice of other mutant lines). If the *shani* mutant phenotype was caused by a single autosomal recessive mutation, *shani* mutant mice were expected to be homozygous for the phenotype-causing mutation and share this mutation. Thus variants were further filtered to only include those that were called as homozygous, had a minimum sequencing depth of five reads, and were not known between-strain SNPs. Next we searched for variants that were shared between all five G3 *shani* mutant mice. This yielded three single nucleotide substitutions at the following positions: Chr11:73187728 in intron of *Ctns* (A to G substitution), Chr11:73336037 in exon of *Spata22* (A to G substitution) and Chr11:76138184 in intron of *Vps53* (A to T substitution).

### ENCODE data analysis

ENCODE long-RNA sequencing data (release 3) for Ensembl Transcript ID: ENSMUST00000092926 with the following GEO accession numbers were used in **Fig. 2d**: testis GSM900193, liver GSM900195, mammary gland GSM900184, large intestine GSM900189, cerebellum GSM1000567, frontal lobe GSM1000562, cortex GSM1000563, kidney GSM900194, thymus GSM900192, heart GSM900199, colon GSM900198, spleen GSM900197, adrenal gland GSM900188, duodenum GSM900187, lung GSM900196, ovary GSM900183, small intestine GSM900186, stomach GSM900185.

### Phylogenetic analysis

SPATA22 protein sequences were obtained from NCBI or Ensembl. Two conserved domains were identified in ten placental mammals using Multalin (Corpet 1988). Sequences of these two domains were used to search for homologous sequences in other species with tBlastn tool (NCBI) and the following databases: Whole Genome Shotgun contigs (WGS), Transcriptome Shotgun Assembly (TSA) and Sequence Read Archive (SRA). Protein sequences obtained were aligned with Clustal Omega (Sievers et al. 2011; Madeira et al. 2019) and edited with Jalview 2.10.5 (Waterhouse et al. 2009). Then conserved domain boundaries were refined as follows: succession of 10 amino acids in which at least half of them possess 40% identity among all listed species. For graphical representation (**Fig. 2e**), consensus sequences were defined for closely related species (10 placentals, 6 marsupials, 4 turtles, 4 snakes, 5 birds, 5 frogs, 4 crocodiles, 12 fishes, 5 echinoderms, 6 cnidarians, 15 molluscs) using Multalin, realigned with Clustal Omega and edited with Jalview. Names of species and corresponding sequence IDs are provided in **Table S1**.

*Spata22* nucleotide sequences were obtained from Ensembl or NCBI. Nucleotide sequence of mouse *Spata22* exon 4 was used to search for homologous sequences with BLASTn tool in Ensembl. Retrieved sequences were aligned with Clustal Omega and edited with Jalview (**Fig. 2f**). Sequences presented correspond to a complete exon and have the following Ensembl exon ID or NCBI gene ID: mouse (ENSMUSE00001408004), rat (ENSRNOE00000381274), human (ENSE00003580316), cow (ENSBTAE00000171295), platypus (ENSOANE00000227841), elephant (ENSLAFE00000151975), tiger (ENSPTIE00000161305), opposum (ENSMODE00000294904), turtle (ENSPSIE00000164161), flycatcher (ENSFALE00000067395) and xenopus (100485180).

### Histology

For secondary screening, frozen testes were embedded in optimal cutting temperature compound (Tissue-Tek, Sakura Finetek, VWR) and sectioned (5 µm) at −20°C. Sections were placed on slides and slides were fixed in 4% paraformaldehyde (PFA) for 15 min at room temperature. Slides were subsequently rinsed in water and stored for up to one week at 4°C prior to staining.

For analyses of *Spata22* mutants, testes and ovaries were fixed overnight in 4% PFA at 4°C, followed by two 5 min washes in water at room temperature and transferred to 70% ethanol. Alternatively, testes were fixed in Bouin’s fixative overnight at room temperature, washed in water for 1 hr at room temperature, followed by two 1 hr washes in 70% ethanol at room temperature. Fixed tissues were stored in 70% ethanol at 4°C for up to one week prior to embedding, embedded in paraffin, and sectioned (5 µm).

Periodic acid-Schiff (PAS) staining and immunohistochemical TUNEL assay were performed by the MSKCC Molecular Cytology Core Facility. PAS staining was done with hematoxylin counterstain using the Autostainer XL automated stainer (Leica Microsystems). TUNEL assay was performed using the Discovery XT processor (Ventana Medical Systems) as previously described (Jain et al. 2017). For immunofluorescent staining, slides were deparaffinized with Xylene, rehydrated with serial washes in 100–70% ethanol and rinsed with water. Antigen retrieval was performed by incubating slides for 10 min at 95– 100°C in 10mM sodium citrate with 0.05% Tween-20 pH 6. Slides were stored at room temperature until staining.

### Chromosome spreads

Spermatocyte spreads were prepared using established methods (Peters et al. 1997; Moens et al. 2000). Briefly, spermatocytes were homogenized in 0.1 M sucrose, then surface spread onto glass slides covered with 1% PFA containing either 0.1% Triton X-100 or 0.25% NP40. Slides were incubated for 2 hr in a humid chamber, gently air-dried and rinsed with 0.4% Photo-Flo (Kodak). Slides were subsequently air-dried and stored at −20°C.

### Immunofluorescence

Chromosomes spreads or testis sections were incubated in blocking buffer (1× PBS with 0.2% BSA, 0.2% cold water fish skin gelatin, 0.05% Tween-20) for 1 hr at room temperature. Slides were incubated with primary antibody overnight at room temperature and washed three times with 1× PBS containing 0.05% Tween-20 at room temperature. Secondary antibody staining was done with appropriate Alexa Fluor secondary antibodies for 1 hr at 37°C and slides were washed three times in 1× PBS containing 0.05% Tween-20 at room temperature. All antibodies were diluted in blocking buffer. Slides were mounted with coverslips using mountant (ProLong Gold, Life Technologies) containing DAPI and stored at 4°C until imaging. Antibodies and dilutions used are listed in **Table S3**.

### Image analysis

PAS-stained and TUNEL slides were digitized using Pannoramic Flash 250 (3DHistech) with 20×/0.8 NA air objective. Images were produced and analyzed using the Pannoramic Viewer software (3DHistech). Immunofluorescence images of testis sections were produced using an SP8 confocal microscope (Leica Microsystems) with 63×/1.4 NA oil-immersion objective. Chromosome spreads were imaged on an AX70 epifluorescent microscope (Olympus) equipped with a CoolSNAP MYO CCD camera (Photometrics) using a 100×/1.35 NA oil-immersion objective.

All histological and cytological data are representative of ≥2 biological replicates (independent mice) of similar ages. Presence/absence of developing follicles in mutant ovaries (**Fig. 3d, e**) was assessed by examining ≥30 sections (spanning ≥150 µm of ovarian tissue). Quantitative data were acquired by manual scoring. For TUNEL quantification (**Fig. 3c**), ≥3 testis sections (≥50 µm apart) were scored per mouse. For detection of number of foci (**Fig. 5, 6)**, immunofluorescence staining was adjusted to be above background level (region containing no cells) using Fiji (Schindelin et al. 2012) and only foci that overlapped with SYCP3 staining (axis-associated foci) were counted. Spermatocytes were staged on the basis of SYCP3 staining patterns. Statistical analyses were done using GraphPad Prism 7.

### Spata22 coding region amplification and sequencing

Procedures involving commercial kits were performed according to manufacturers’ instructions. Total RNA was extracted from frozen testes using the RNeasy Mini Kit (Qiagen) and the RNase-Free DNase Set (Qiagen) was used to perform an on-column DNase digestion. cDNA was synthesized from 1 µg of RNA using High-Capacity cDNA Reverse Transcription Kit (Applied Biosystems) with random primers.

*Spata22* coding region was amplified from adult testis cDNA by PCR with *Spata22* coding sequence amplification F and R primers using Phusion High Fidelity DNA Polymerase (Thermo Fisher Scientific) and 0.3 ng/µl of cDNA in GC Buffer. Manufacturer’s thermocycling conditions were adapted as follows: denaturation for 10 sec, annealing for 30 sec at 65°C and extension for one min for a total of 35 cycles. PCR products were electrophoresed on a 0.8% agarose gel, bands corresponding in size to *Spata22* cDNA were gel purified using QIAquick PCR Purification Kit (Qiagen) and Sanger sequenced using *Spata22* seq1–seq5 primers. Primer sequences are provided in **Table S2**.

### Quantitative RT-PCR

Testis cDNA was obtained as described above. Procedures involving commercial kits were performed according to manufacturers’ instructions. Quantitative PCR was carried out using QuantiFast SYBR Green PCR Kit (Qiagen) for detecting products on a StepOnePlus Real-Time PCR instrument (Applied Biosystems). Primer sequences are listed in **Table S2**.

All reactions were done in duplicate (adult mice) or quadruplicate (12 dpp-old mice). The success of reactions was confirmed by analysis of amplification and melting curves. StepOne Software v2.3 was used to quantify products using the 2^-ΔΔCt^ method. Briefly, duplicate or quadruplicate threshold cycle (Ct) values were averaged to obtain mean Ct values and mean Ct value of *Actb* reactions was subtracted to obtain ΔCt values. Then ΔCt values obtained for mutants were subtracted from those obtained for wild-type or heterozygous littermates to obtain ΔΔCt values and 2^-ΔΔCt^ values are plotted as relative quantity values (**Fig. 7c**).

### Plasmid construction

Constructs were confirmed by DNA sequencing. The *shani* mutation was introduced into pCMV6-XL5-HsSPATA22 plasmid (Origene, cat# SC123038) by site directed mutagenesis. Wild-type and mutated *Spata22* sequences were PCR amplified from pCMV6-XL5-HsSPATA22, digested with EcoRI (NEB) and BamHI (NEB), and cloned into pKH3 plasmid (Addgene) for protein expression.

### Cell culture and plasmid transfection

Cells were cultivated in a 5% CO_2_ humidified incubator at 37°C. Human embryonic kidney cells (HEK-293, ATCC) were grown in Dulbecco’s modified Eagle medium supplemented with 10% fetal bovine serum (Gibco). Cell lines were regularly screened for mycoplasma contamination by DAPI staining. Plasmid transfection was performed with Lipofectamine 2000 (Life Technologies) according to the manufacturer’s instructions.

### Western blot analysis

Total protein extraction from testes was done, after removing the tunica albuginea, by lysing cells in RIPA buffer (Thermo Fisher Scientific) using ceramic spheres and FastPrep-24 instrument (MP Biomedicals) with two 20 sec pulses. Extracts were then sonicated for 10 min at medium power (30 sec on, 30 sec off) and centrifuged at 16,200 ×g for 10 min at 4°C. For whole cell protein extraction from HEK-293 cells, cells were lysed under native conditions by homogenization in 1× cell lysis buffer (Cell Signaling) supplemented with Complete protease inhibitor tablet (Roche), 1 mM phenylmethylsulfonyl fluoride (Sigma) and 10 µM MG-132 (Sigma), sonicated twice for 10 min each at medium power (30 sec on, 30 sec off) and centrifuged at 16,200 ×g for 10 min at 4°C.

Protein samples were supplemented with Laemmli buffer and dithiothreitol, boiled for five min and separated by SDS-PAGE on a 10% gel. Proteins were transferred onto a polyvinylidene difluoride membrane. Membranes were blocked in PBST (1× PBS with 0.05% Tween-20) supplemented with 5% non-fat milk for 1 hr at room temperature, incubated with primary antibodies in PBST supplemented with 1% non-fat milk overnight at 4°C and then incubated with appropriate secondary antibodies in PBST supplemented with 1% non-fat milk for 3 hr at room temperature. Images were acquired using a Typhoon 5 imager (Amersham Biosciences, GE Healthcare). Antibodies used are listed in **Table S3**.

## End Matter

### Author Contributions and Notes

CP, VB, JR, EM and DJ performed research. CP, NL, GL, EM and DJ analyzed data. GL, SK, EM and DJ wrote the paper. The authors declare that they have no conflict of interest. Reagents and mouse strains are available upon request. All applicable international, national, and/or institutional guidelines for the care and use of animals were followed. All procedures performed in studies involving animals were in accordance with the regulatory standards and were approved by the Memorial Sloan Kettering Cancer Center (MSKCC) and the Commissariat à l’énergie atomique et aux énergies alternatives (CEA) Institutional Animal Care and Use Committees.

## Acknowledgments

We thank Keeney lab members Luis Torres, Diana Y. Eng, and Jacquelyn Song for assistance with genotyping and mouse husbandry. We thank Keeney lab member Shintaro Yamada for *Spo11* mutant mice. We thank Ning Fan and Dmitry Yarilin at the MSKCC Cytology Core for help with histological analyses. For whole-exome sequencing, we thank the MSKCC Integrated Genomics Operation. We thank members of the Laboratory of the Development of the Gonads as well as Keeney lab members for discussions and technical support. We thank members of the animal housing facility at CEA-FAR and CEA-Saclay. We acknowledge the ENCODE Consortium (Consortium 2012) and the ENCODE production laboratory of Thomas Gingeras (Cold Spring Harbor Laboratory) for generating the ENCODE datasets used in this manuscript.

MSKCC core facilities were supported by National Institutes of Health grant P30 CA008748. Livera lab members were supported by ANR, INSERM, Fondation ARC and La Ligue contre le Cancer. DJ was supported by a fellowship from Human Frontier Science Program. Work in the SK lab was supported by Howard Hughes Medical Institute. CP and JR were supported by a CEA fellowship and JR by La Ligue contre le Cancer.

---

**Table S1.** Accession numbers of sequences used for phylogenetic analysis.

| **Group** | **Species** | **ID** |
| --- | --- | --- |
| Eutheria | *Mus musculus* | NP_001038996.1 |
|  | *Rattus norvegicus* | NP_001178753.1 |
|  | *Homo sapiens* | NP_001164169.1 |
|  | *Macaca fascicularis* | NP_001270464.1 |
|  | *Oryctolagus cuniculus* | XP_002718916.1 |
|  | *Capra hircus* | XP_005693459.2 |
|  | *Panthera tigris altaica* | XP_007076209.1 |
|  | *Eptesicus fuscus* | XP_008154995.1 |
|  | *Loxodonta Africana* | XP_003417051.1 |
|  | *Trichechus manatus latirostris* | XP_004376192.1 |
| Metatheria | *Phascolarctos cinereus* | ENSPCIP00000026657.1 |
|  | *Notamacropus eugenii* | WGS and SRA |
|  | *Vombatus ursinus* | WGS |
|  | *Monodelphis domestica* | XP_007506518.1 |
|  | *Sarcophilus harrisii* | XP_023357058.1 |
|  | *Perameles nasuta* | TSA |
| Testudines | *Terrapene mexicana triunguis* | XP_024067042.1 |
|  | *Chrysemys picta bellii* | XP_023961688.1 |
|  | *Pelodiscus sinensis* | XP_025035057.1 |
|  | *Malaclemys terrapin* | WGS |
| Aves | *Struthio camelus australis* | KFV80192.1 |
|  | *Pygoscelis adeliae* | KFW70311.1 |
|  | *Gavia stellata* | KFV51333.1 |
|  | *Buceros rhinoceros silvestris* | KFO88856.1 |
|  | *Apteryx rowi* | XP_025911051.1 |
| Crocodilia | *Alligator mississippiensis* | XP_014459223.1 and SRA |
|  | *Alligator sinensis* | XP_014379890.1 |
|  | *Crocodylus porosus* | XP_019401285.1 |
|  | *Gavialis gangeticus* | XP_019366723.1 |
| Serpentes | *Pseudonaja textilis* | XP_026566041.1 |
|  | *Notechis scutatus* | XP_026544297.1 |
|  | *Protobothrops mucrosquamatus* | XP_015681783.1 |
|  | *Python bivittatus* | XP_025029367.1 |
| Anura | *Xenopus tropicalis* | XP_012812971.1 |
|  | *Nanorana parkeri* | XP_018413137.1 |
|  | *Rhinella marina* | WGS and SRA |
|  | *Bufo viridis* | TSA |
|  | *Odorrana tormota* | TSA |
| Actinopterygii | *Astyanax mexicanus* | XP_007260432.2 |
|  | *Danio rerio* | XP_021332340.1 |
|  | *Haplochromis burtoni* | TSA |
|  | *Oreochromis niloticus* | TSA |
|  | *Acipenser oxyrinchus* | TSA |
|  | *Anguilla anguilla* | TSA |
|  | *Salmo salar* | TSA |
|  | *Esox lucius* | XP_010869021.1 |
|  | *Xiphophorus maculatus* | XP_014324914.1 |
|  | *Electrophorus electricus* | XP_026859897.1 |
|  | *Lepisosteus oculatus* | XP_015222922.1 |
|  | *Hippocampus comes* | XP_019750777.1 |
| Echinodermata | *Patiria pectinifera* | TSA |
|  | *Acanthaster planci* | XP_022093358.1 |
|  | *Strongylocentrotus purpuratus* | XP_011682669.1 |
|  | *Apostichopus japonicus* | PIK60563.1 |
|  | *Evechinus chloroticus* | TSA |
| Cnidaria | *Pocillopora damicornis* | XP_027057005.1 |
|  | *Stylophora pistillata* | XP_022794271.1 |
|  | *Orbicella faveolata* | XP_020616570.1 |
|  | *Favia lizardensis* | TSA |
|  | *Nematostella vectensis* | XP_001633929.1 |
|  | *Hydra vulgaris* | XP_012561417.1 |
| Mollusca | *Elysia chlorotica* | RUS76875.1 |
|  | *Lottia gigantea* | XP_009051722.1 |
|  | *Aplysia californica* | XP_005102185.1 |
|  | *Biomphalaria glabrata* | XP_013065423.1 |
|  | *Biomphalaria pfeifferi* | TSA |
|  | *Limacina retroversa* | TSA |
|  | *Elysia timida* | TSA |
|  | *Lymnaea stagnalis* | TSA |
|  | *Crassostrea virginica* | XP_022302499.1 |
|  | *Mizuhopecten yessoensis* | XP_021365932.1 |
|  | *Nodipecten subnodosus* | TSA |
|  | *Mytilus galloprovincialis* | TSA |
|  | *Adamussium colbecki* | TSA |
|  | *Scapharca broughtonii* | TSA |
|  | *Octopus bimaculoides* | KOF91762.1 |
| Porifera | *Amphimedon queenslandica* | XP_019848728.1 |
| Arthropoda | *Drosophila melanogaster* | NP_001246272.1 |
|  | *Plutella xylostella* | TSA |
|  | *Zootermopsis nevadensis* | XP_021921944.1 |
|  | *Acyrtosiphon pisum* | XP_016661503.1 |

**Table S2.**
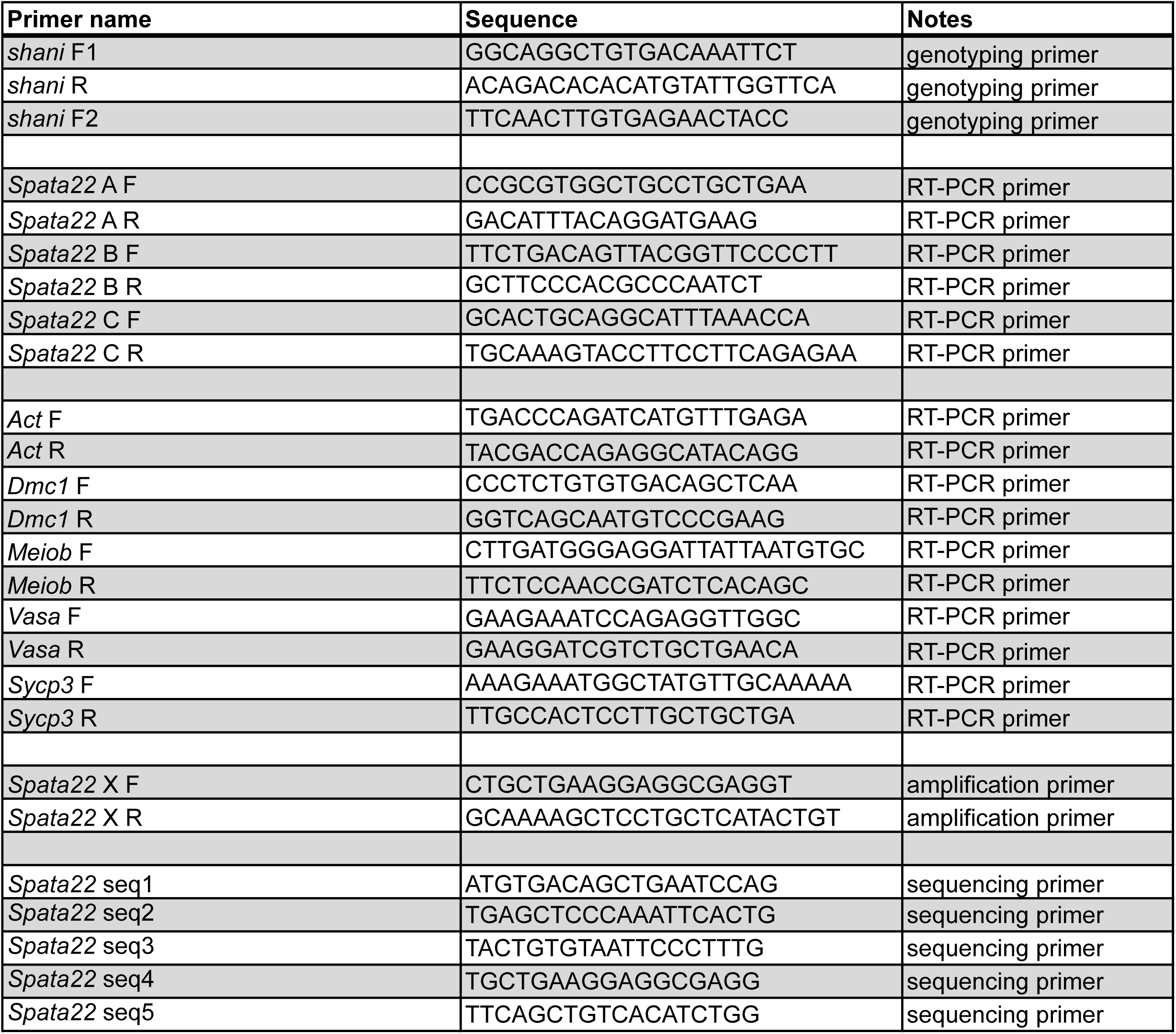
Oligonucleotide primers.

**Table S3.** Antibodies.

| **Target** | **Species** | **Catalog/Reference number** | **Source** | **Dilution for immunofluorescence** | **Dilution for immunoblotting** |
| --- | --- | --- | --- | --- | --- |
| RPA2 | Rat | 2208 | Cell Signaling | 1:400 | - |
| HA tag | Rabbit | Ab9110 | Abcam | - | 1:1000 |
| MEIOB | Rabbit | AK2 | Livera's lab | 1:200 | - |
| SPATA22 | Rabbit | 16989-1-AP | Proteintech | 1:400 | 1:1000 |
| DMC1 | Rabbit | SC-22768 | Santa Cruz Biotechnology | 1:200 | - |
| SYCP3 | Mouse | Ab7672 | Abcam | 1:500 | - |
| SYCP3 | Rabbit | NB300-232 | Novus Biologicals | 1:500 | - |
| SYCP1 | Rabbit | Ab15090 | Abcam | 1:500 | - |
| BLM | Rabbit | NB100-214 | Novus Biologicals | 1:200 | - |
| γH2AX | Mouse | 05-636 | Millipore | 1:500 | - |
| LAMIN B1 | Mouse | SC-374015 | Santa Cruz Biotechnology | - | 1:500 |
| Anti-mouse, Alexa Fluor 488 | Donkey | 715-545-151 | Jackson ImmunoResearch | 1:500 | - |
| Anti-mouse, Alexa Fluor 594 | Donkey | 715-585-151 | Jackson ImmunoResearch | 1:500 | 1:1000 |
| Anti-rabbit, Alexa Fluor 594 | Donkey | 711-585-152 | Jackson ImmunoResearch | 1:500 | 1:1000 |
| Anti-rabbit, Alexa Fluor 488 | Donkey | 711-545-152 | Jackson ImmunoResearch | 1:500 | - |
| Anti-rat, Alexa Fluor 647 | Donkey | 712-605-153 | Jackson ImmunoResearch | 1:500 | - |
| Anti-rat, Alexa Fluor 488 | Donkey | 712-545-153 | Jackson ImmunoResearch | 1:500 | - |

